# Implications for tetraspanin-enriched microdomain assembly based on structures of CD9 with EWI-F

**DOI:** 10.1101/2020.06.02.130047

**Authors:** Wout Oosterheert, Katerina T. Xenaki, Viviana Neviani, Wouter Pos, Sofia Doulkeridou, Jip Manshande, Nicholas M. Pearce, Loes M. J. Kroon-Batenburg, Martin Lutz, Paul M. P. van Bergen en Henegouwen, Piet Gros

## Abstract

Tetraspanins are ubiquitous eukaryotic membrane proteins that contribute to a variety of signaling processes by spatially organizing partner-receptor molecules in the plasma membrane. How tetraspanins bind and cluster partner receptors into so-called tetraspanin-enriched microdomains is unknown. Here we present crystal structures of the large extracellular loop of CD9 in complex with nanobodies 4C8 and 4E8; and, the cryo-EM structure of 4C8-bound CD9 in complex with its prototypical partner EWI-F. The CD9 - EWI-F complex displays a tetrameric arrangement with two centrally positioned EWI-F molecules, dimerized through their ectodomains, and two CD9 molecules, one bound to each EWI-F single-pass transmembrane helix through CD9-helices h3 and h4. In the crystal structures, nanobodies 4C8 and 4E8 bind CD9 at the C and D loop, in agreement with 4C8 binding at the ends of the CD9 - EWI-F cryo-EM complex. Overall, the 4C8 - CD9 - EWI-F - EWI-F - CD9 - 4C8 complexes varied from nearly two-fold symmetric (i.e. with the two CD9 - 4C8 copies in nearly anti-parallel orientation) to ca. 50° bent arrangements. Since membrane helices h1 and h2 and the EC2 D-loop have been previously identified as sites for tetraspanin homo-dimerization, the observed linear but flexible arrangement of CD9 - EWI-F with potential CD9 - CD9 homo-dimerization at either end provides a new ‘concatenation model’ for forming short linear or circular assemblies, which may explain the occurrence of tetraspanin-enriched microdomains.

## Introduction

The spatial organization of proteins and lipids in the plasma membrane controls cellular processes such as signaling, trafficking, cell adhesion and fusion (1). Members of the tetraspanin superfamily of transmembrane proteins act as molecular organizers of the plasma membrane by clustering specific partner proteins in *cis* to arrange their assembly required for function (2–5). Tetraspanins form dynamic platforms termed tetraspanin-enriched microdomains (TEMs), also referred to as the tetraspanin web (3, 4, 6). TEMs were formerly described as large assemblies comprising multiple tetraspanin family members and partner proteins (6–8), but the most recent model derived from super-resolution microscopy experiments states that TEMs are nanoclusters of approximately 120 nm in diameter that contain fewer than ten copies of a single tetraspanin homolog (9).

Among the 33 human tetraspanins, CD9 (also known as Tspan29) is an extensively studied tetraspanin that is expressed in a wide variety of tissues and is also highly abundant on the membranes of extracellular vesicles (10). CD9 participates in processes like cell adhesion, motility and differentiation (4, 11, 12). Additionally, it regulates cell-cell fusion events in myoblasts (13, 14), osteoclasts (15), macrophages (16) and sperm-egg cells (17, 18). Besides its physiological functions, CD9 is widely associated with diseases: both the upregulation and downregulation of CD9 are linked with poor prognosis in several types of cancer, such as melanoma, leukemia and gastric, lung, breast, colon and prostate malignancies (19). Moreover, CD9 modulates HIV-1-induced membrane fusion (20) and promotes MERS-coronavirus entry by clustering host-cell receptors and proteases (21, 22).

At the molecular level, tetraspanins adopt a common architecture with four transmembrane helices (h1 - h4), intracellular N- and C-termini, a small extracellular loop between membrane helices h1 and h2 (EC1) and a large extracellular loop between membrane helices h3 and h4 (EC2). Initial structural studies on the soluble EC2 domain of CD81 disclosed an EC2-fold with five helical regions (termed A - E), with A, B and E forming a ‘stalk’ domain and C and D forming a ‘head’ domain (23). Crystal structures of full-length tetraspanin CD81 (24) and, recently, CD9 with a truncated EC2 (25), revealed an inverted-cone shape arrangement of the transmembrane domain (TMD), with the EC2 closing off the TMD-cone like a lid, although molecular dynamics simulations suggested that the EC2 lid can also adopt open conformations (24, 25). Previous studies have established that both the EC2 and TMD regions of tetraspanins participate in homo- and hetero-oligomeric interactions, indicating that tetraspanins employ separate domain regions to bind to different partners proteins (5). CD9 interacts with numerous single-span transmembrane proteins, including integrins (26–29), immunoglobin superfamily (IgSF) proteins (30–32), heparin-binding EGF-like growth factor (33) and metalloprotease ADAM17 (11, 34). IgSF protein EWI-F, also known as CD9-partner 1 or prostaglandin F2 receptor negative regulator, is a prototypical primary CD9-binding protein that also interacts with its close-homolog CD81 in stoichiometric amounts in several cell types (30, 32). EWI-F comprises six extracellular, heavily glycosylated Ig-like C2 domains (35), a single transmembrane (TM) helix and a small C-terminal cytoplasmic tail. Functionally, EWI-F moderates CD9 and CD81-driven cellular fusion events (14, 36) and connects TEMs to the cytoskeleton by binding to ezrin-radixin-moesin proteins (37). Biochemical experiments have revealed that the interaction between EWI-F and CD9 or CD81 is mediated by the TMDs, with critical roles for the transmembrane helix of EWI-F and helix h4 of CD9 or CD81 (36, 38). Additionally, we recently found that palmitoylated membrane-proximal cysteines of CD9 are crucial for maintaining the interaction with EWI-F (39).

Although tetraspanins have a central role in human physiology, the mechanisms through which they interact with partner proteins and the molecular principles that govern the formation and signaling of TEMs remain incompletely understood. Here, we study the structural basis for interactions in the tetraspanin web using the CD9 - EWI-F complex as a model system. We first present crystal structures of the isolated EC2 domain of CD9 in the absence and presence of anti-CD9 nanobodies 4C8 and 4E8, revealing the flexibility of the D-loop region of CD9. We then employ nanobody 4C8 in single-particle cryo-EM studies to show that CD9 and EWI-F form a hetero-tetramer that adopts a range of conformations, with a central EWI-F dimer flanked by two CD9 molecules on each side. The structural data are in good agreement with the recently reported cryo-EM map of CD9 bound to EWI-F homolog EWI-2 (25), indicating that CD9 binds to EWI proteins through a common binding mode. Overall, the various conformations of the CD9 - EWI-F complex observed in the cryo-EM data, combined with previous biochemical interaction studies, support a ‘concatenation model’ for the assembly of TEMs in the plasma membrane.

## Results

### Crystal structures of CD9_EC2_ in complex with nanobodies

Our group has previously reported the generation of several nanobodies that bind to the extracellular region of CD9 (39). We selected nanobodies 4C8 and 4E8, which differ in amino-acid composition in complementarity-determining regions (CDRs) 2 and 3, for further characterization. 4C8 and 4E8 bind to purified, full-length CD9 with affinities of 0.9 nM and 2.0 nM, respectively, and to endogenous CD9 expressed on HeLa cells with affinities of 0.6 nM and 33 nM, respectively (Fig. S1A, B). To determine the nanobody-binding epitopes on CD9, we solved crystal structures of the soluble EC2 of CD9 (CD9_EC2_) in complex with 4C8 at 2.7-Å resolution (Fig. 1A) and 4E8 at 1.4-Å resolution (Fig. 1B), as well as the structure of 4C8 alone at 1.7-Å resolution (Fig. S2A, Table 1). The CD9_EC2_ - 4C8 complex crystallized in space group *I2_1_2_1_2_1_* with two copies in the asymmetric unit (Fig. S2B), whereas CD9_EC2_ - 4E8 crystallized in space group *P2_1_2_1_2* with a single copy of the complex in the asymmetric unit (Fig. 1B).

**Figure 1:**
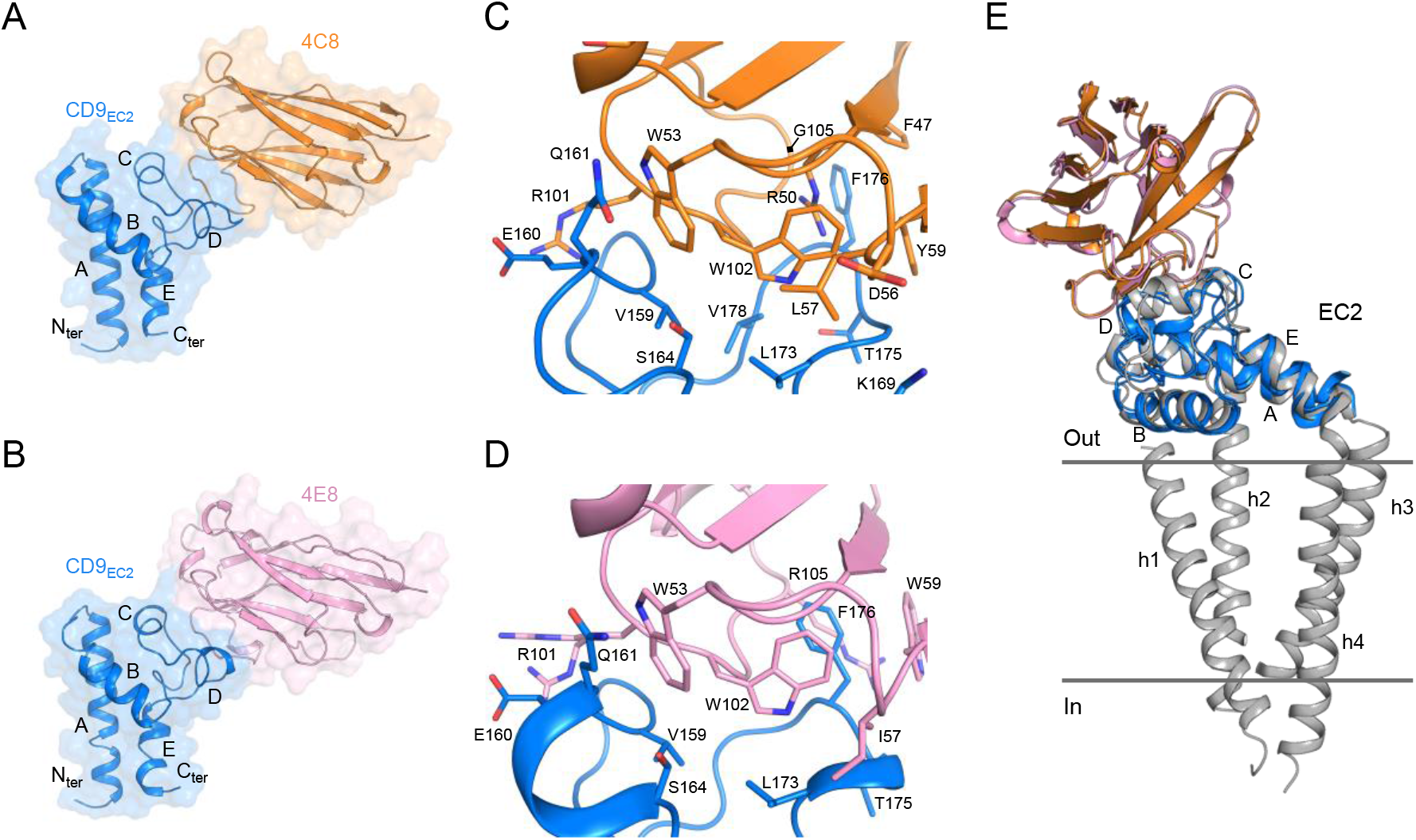
Nanobody-bound CD9_EC2_ structures. (A, B) Crystal structure of CD9_EC2_ (blue) bound to nanobody 4C8 (orange, panel A) and nanobody 4E8 (pink, panel B) The regions of the EC2 are annotated. (C, D) Interaction interface between CD9_EC2_ (blue) and nanobody 4C8 (orange, panel C) and nanobody 4E8 (pink, panel D). Residues contributing to the interface are shown as sticks. (E) Overlay of the nanobody-bound EC2 structures with the structure of full-length CD81 (grey). The CD81 structure is shown parallel to the membrane as a sideview.

**Table 1:**
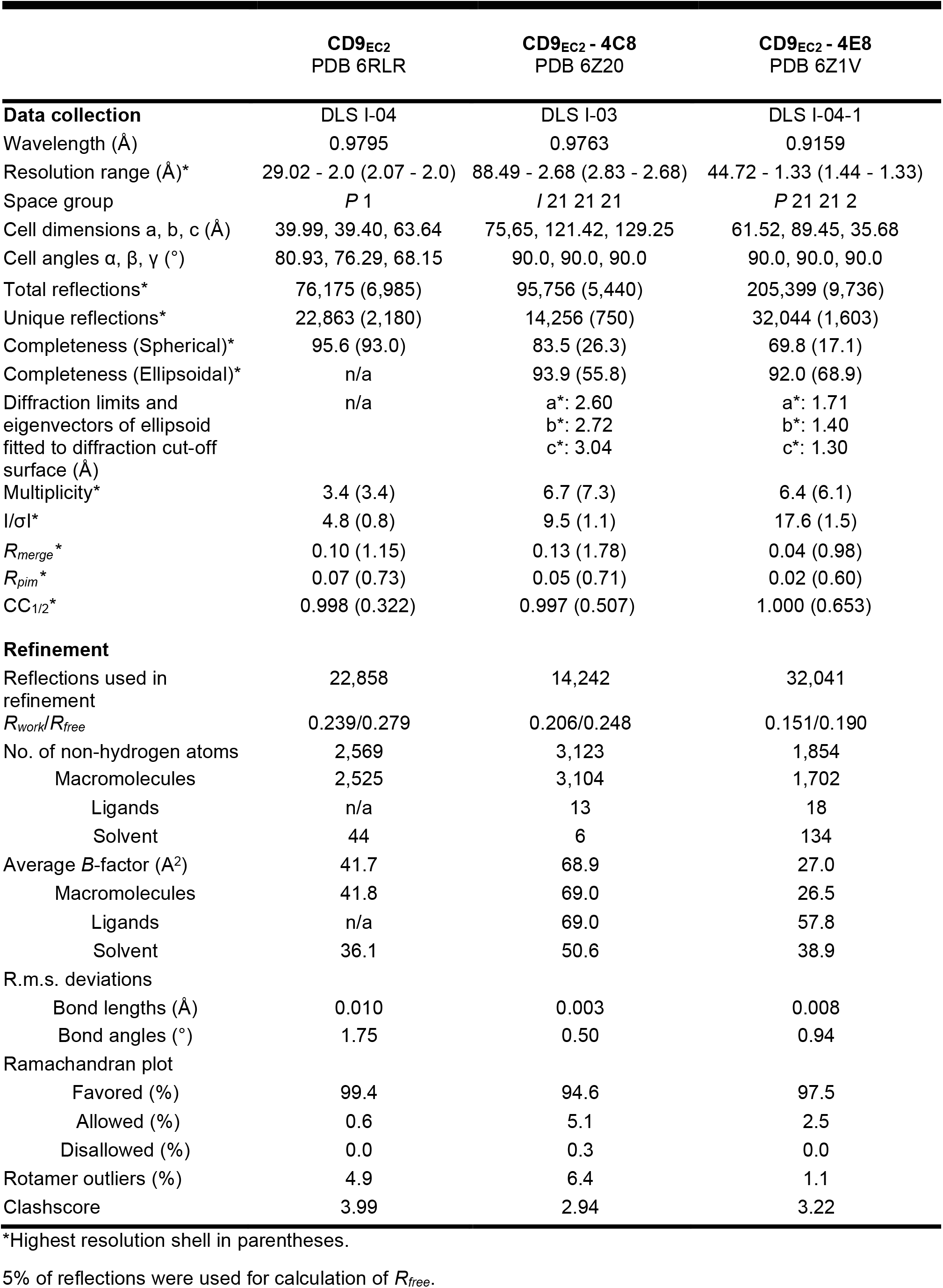
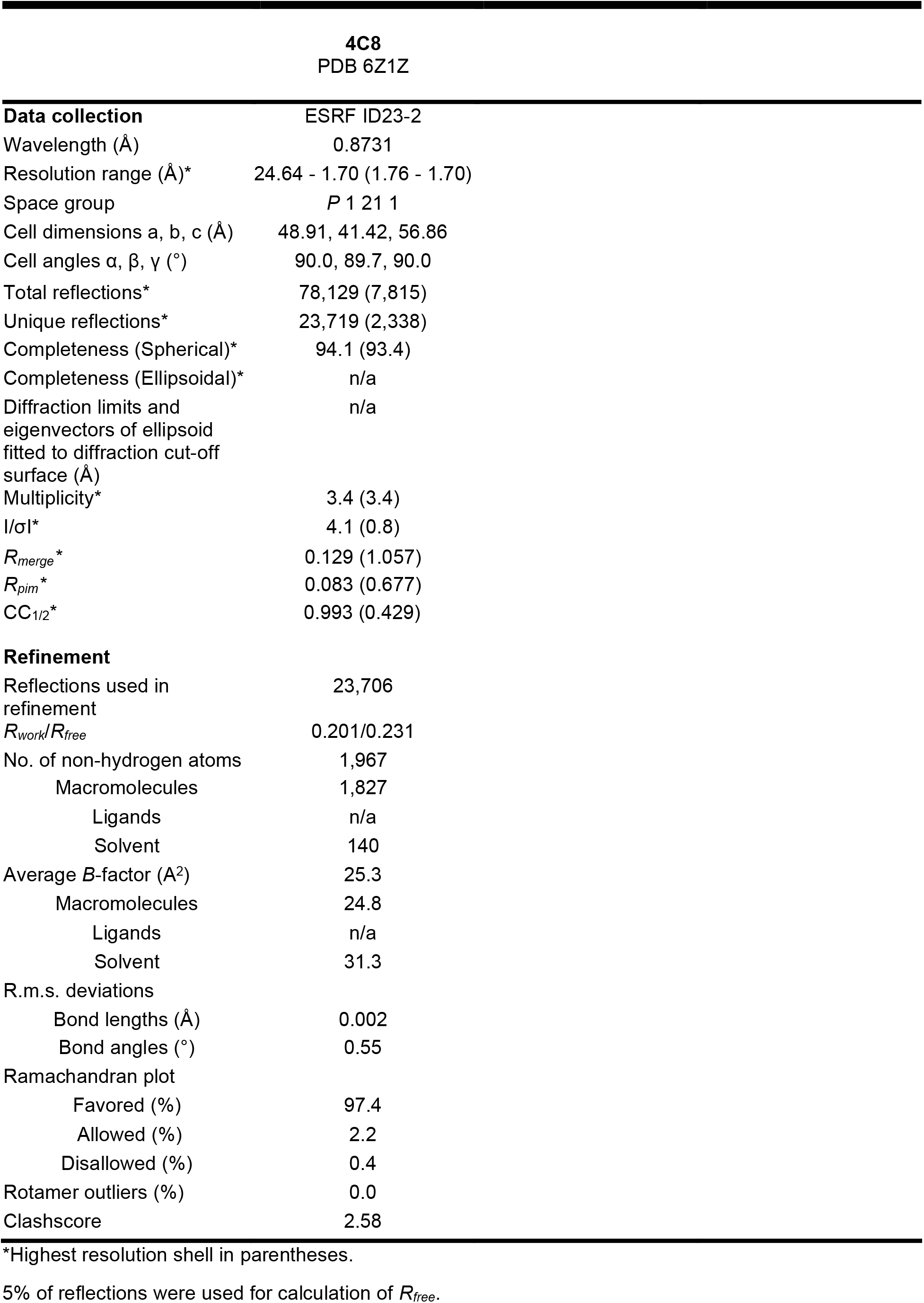
Crystallographic and structure refinement statistics.

In both nanobody-bound structures, CD9_EC2_ adopts an arrangement that is globally similar to previously reported EC2-structures of CD81 (23, 24, 40–43). 4C8 and 4E8 bind nearly identical epitopes that span the C and D loops of CD9_EC2_ (Fig. 1A, B). The interfaces are established through nanobody residues located in CDRs 2 and 3, whereas the CDR1 region of both nanobodies makes no contacts with CD9_EC2_. The surface areas buried in the CD9_EC2_ - 4C8 and 4E8 complexes are ca. 1,280-Å^2^ and 1,230-Å^2^, respectively. CD9 residues involved in the interfaces with the nanobodies are V159, E160, Q161 and S164 from the C loop; and K169, L173, T175, F176 and V178 from the D loop (Fig. 2C, D). Both nanobodies feature an arginine residue (R101) in CDR3 that interacts with E160 through a salt bridge, and two tryptophans (W53 of CDR2 and W102 of CDR3 in both 4C8 and 4E8) that form a hydrophobic core in the interface with CD9_EC2_ (Fig. 1C, D). In CD9_EC2_ - 4C8, the D loop adopts a partially helical conformation and central residue F176 is sandwiched by 4E8 residues W59 of CDR2 and W102 and R105 of CDR3 (Fig. 1D). In the 4C8-bound CD9_EC2_ structure the tip of the D loop points more outward and the Cα atom of F176 is shifted by 3 Å, such that F176 resides not at the center of the interface but at the periphery, where it forms a van der Waals interaction with 4C8-residue G105 (Fig. 1C).

**Figure 2:**
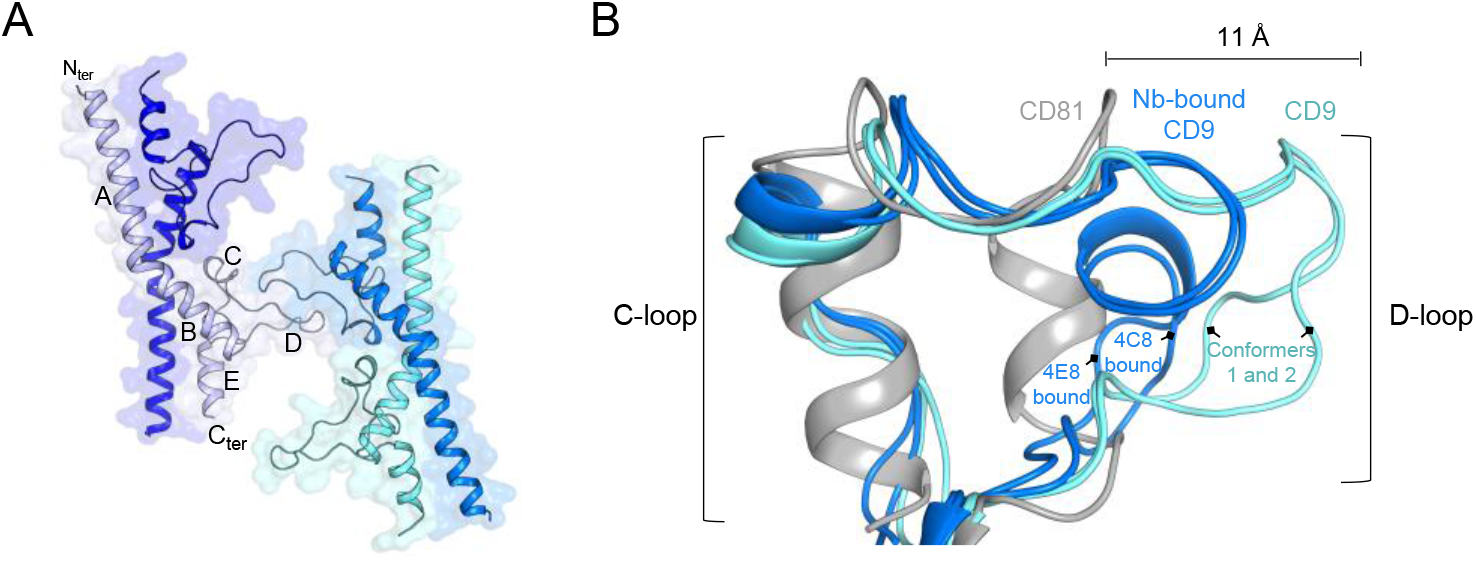
CD9_EC2_ structure and D-loop flexibility. 150:115(A) Asymmetric unit of the twinned CD9_EC2_ crystal, colored by protein chain. The regions of a single EC2 chain are annotated. (B) Overlay of the C- and D-loop arrangements of CD9_EC2_ (cyan), nanobody (Nb) 4C8 and 4E8 bound CD9_EC2_ (blue) and full-length CD81 (grey, pdb 5TCX).

A superimposition of the nanobody-bound CD9_EC2_ structures onto the crystal structure of full-length CD81 (pdb 5TCX) indicates that 4C8 and 4E8 orient away from the membrane plane in the EC2 conformation adopted by CD81 (Fig. 1E), which is consistent with both nanobodies binding to cell-membrane embedded CD9 (Fig. S1A).

### Conformational flexibility of the CD9_EC2_ D-loop

The EC2-D loop displays a high amino-acid sequence variability among tetraspanins and has been proposed to mediate homo- and hetero-oligomeric interactions (5, 44–46). Previously reported structures of CD81 revealed the conformational plasticity of its EC2 D-loop, with both fully helical and partially unfolded arrangements (23, 24, 40–43). Our nanobody-bound CD9_EC2_ structures also display a minor conformational difference in D-loop with respect to each other (Fig. 1C, D). To further investigate the conformational changes adopted by the D-loop of CD9 upon nanobody binding, we determined the crystal structure of CD9_EC2_ in the absence of nanobodies at 2.0-Å resolution (Table 1). The diffraction data indicated non-merohedral twinning of the crystal with a twofold rotation around the ***a***\*+***b***\* diagonal as the twinning operation (Fig. S3). Moreover, we observed diffuse streaks in the ***a***\*+***b***\* direction (Fig. S3A). Figure S3B shows the two twin domains in the crystal with the twinning interface in the middle. For structure refinement the data was de-twinned based on the calculated structure factors (see Experimental procedures).

CD9_EC2_ crystallized in space group *P1* with four molecules in the asymmetric unit, arranged as a dimer of domain-swapped dimers (Fig. 2A). In this domain swap, the N-terminal A helix (residues 114-138) is exchanged with the A helix of its dimeric partner, resulting in an extensive interface that buries ca. 3,400-Å^2^ of surface area. The two connected protein cores each strongly resemble the monomeric form of CD9_EC2_, which is commonly observed for structures of domain-swapped dimers (47, 48). The domain-swapped dimers are packed in the crystal lattice through interactions between the D-loop regions, which are extended and fully unfolded (Fig. 2A). A comparison between all CD9_EC2_ molecules in the asymmetric unit indicated that the D-loop adopts two distinct conformations, both different from the nanobody-bound CD9_EC2_ structures (Fig. 2B). This observation suggests that the binding of nanobodies 4C8 and 4E8 to CD9_EC2_ changes the D-loop conformation. This may, at least in part, explain the difference in signal intensity between 4C8 and 4E8 binding to purified CD9 and to CD9 on cells (Fig. S1A, B), assuming that the particular conformational change in the D-loop of CD9 is less efficient in the case of 4E8.

An overlay of the C- and D-loop regions of CD9_EC2_ structures in the absence and presence of nanobodies and full-length CD81 (pdb 5TCX) revealed a diverse set of D-loop conformations: the D-loop of CD9_EC2_ is extended ~11 Å compared to the compact helical arrangement observed in the structure of full-length CD81 (Fig. 2B). In the twinned CD9_EC2_ structure, one of the D-loops has large B-factors (Fig. S3C), indicating its flexibility, and it is involved in the twinning interface, apparently accommodating the deviation of true 2-fold symmetry of the two independent molecules. Additionally, the C-loops of the CD9_EC2_ structures adopt a more loop-like conformation compared to the helical C-loops present in CD81 structures (Fig. 2B).

### Purified CD9-EWI-F complex is flexible

We next focused on obtaining a structural model of full-length CD9 in complex with its primary partner EWI-F. To this end, we co-expressed Strep-GFP-tagged CD9 together with EWI-F in HEK293 cells and isolated the complex in N-dodecyl-ß-D-maltoside (DDM) detergent using Strep-affinity chromatography and size-exclusion chromatography (SEC) purification steps (Fig. 3A, B). The complex did not dissociate during subsequent analytical fluorescence assisted SEC experiments (Fig. 3C), indicating that the interaction between CD9 and EWI-F is preserved outside the native cell-membrane environment. We then collected a preliminary single-particle cryo-EM dataset of the purified complex. The resulting micrographs showed elongated, highly heterogeneous protein particles (Fig. 3D). This heterogeneity is likely caused by flexibility between the six Ig-like C2 domains of EWI-F, indicating that the imaged CD9 - EWI-F complex in its full-length version is unsuitable for high-resolution structure determination. 2D-classification experiments yielded low-resolution class averages with no distinguishable micelle region present. Nevertheless, a single 2D-class average showed two rows of five domain-like densities stacked together (Fig. 3E, boxed in red), presumably corresponding to five of the six Ig-like domains of EWI-F. This suggests that EWI-F forms dimeric assemblies, which is consistent with a previous cross-linking study (38).

**Figure 3:**
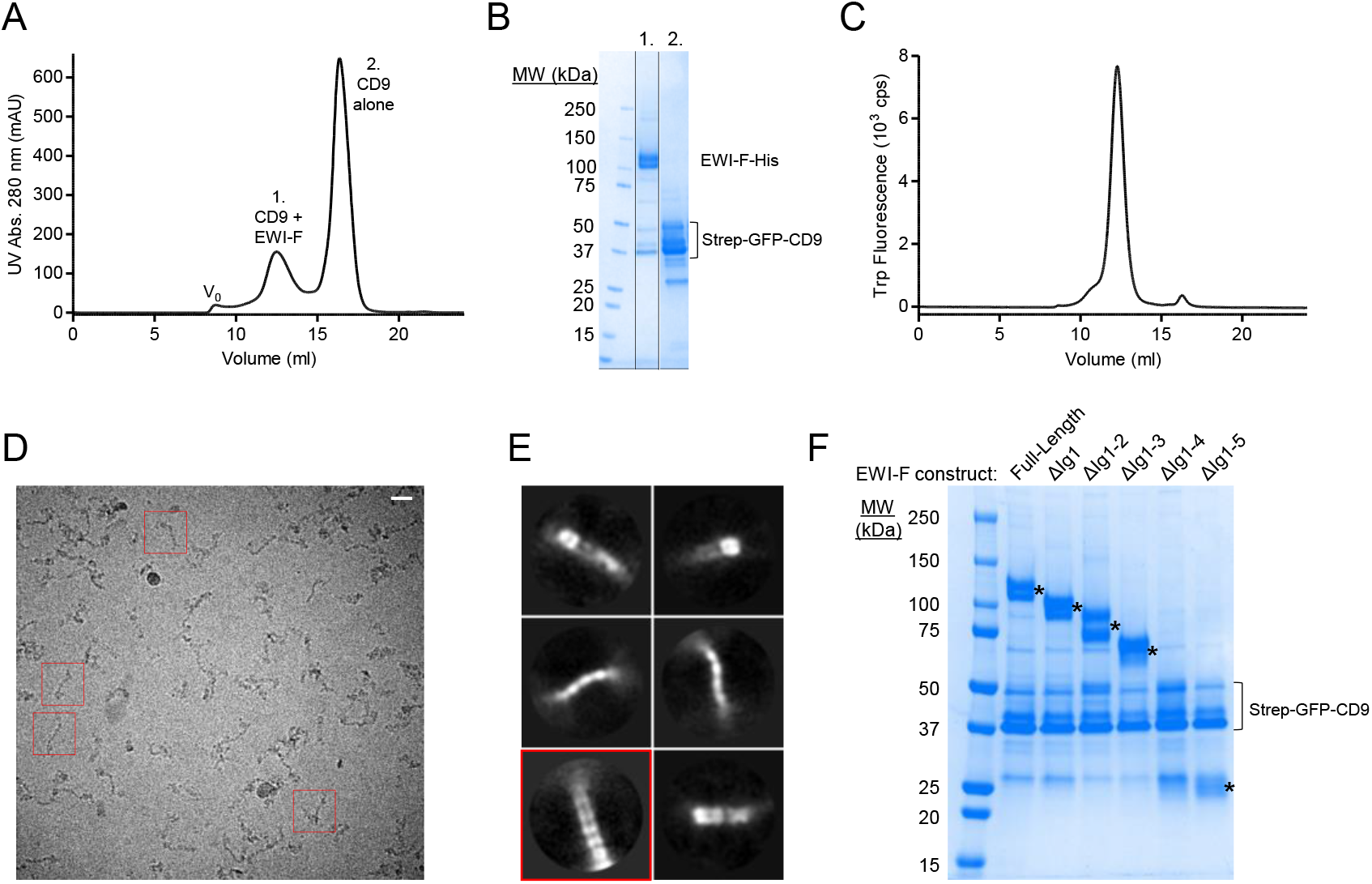
Biochemical and structural characterization of CD9 with full-length EWI-F and EWI-F truncations. (A) Size-exclusion chromatography elution profile of co-expressed CD9 and EWI-F after Strep-affinity purification. (B) SDS page gel of peaks 1 and 2 of the SEC elution from panel A. Multiple bands are visible for EWI-F due to heterogeneous glycosylation, and for CD9 due to the partial SDS-induced unfolding of the GFP-tag. (C) Analytical Tryptophan fluorescence-assisted SEC elution profile of the CD9-EWI-F complex used for cryo-EM. (D) Cryo-EM micrograph depicting CD9 - EWI-F particles in vitreous ice. The scale bar length is 200 Å. Examples of individual particles are boxed in red. (E) Selected 2D-class averages generated through Relion. The box size is 409×409 Å. The class that shows evidence for dimeric EWI-F is boxed in red. (F) SDS page gel of Strep-purified CD9 with different variants of EWI-F. Gel bands at the expected molecular weights of the EWI-F variants are marked with an asterisk (*).

### CD9 interacts with truncated EWI-F variants

Because full-length EWI-F displayed severe inter-domain flexibility, we next aimed to create a structurally more homogeneous EWI-F protein sample. For this purpose, we designed and generated N-terminally truncated EWI-F constructs with up to five Ig-like domains removed, which we termed EWI-F_ΔIg1_, EWI-F_ΔIg1-2_, EWI-F_ΔIg1-3_, EWI-F_ΔIg1-4_ and EWI-F_ΔIg1-5_. To assess if the truncated EWI-F constructs still associated with CD9, we expressed the EWI-F variants together with Strep-GFP-tagged CD9 in small scale HEK293-cell cultures and, after Strep-affinity purification, monitored the amount of co-purified EWI-F using SDS-PAGE. The SDS-PAGE gel revealed the presence of protein bands at expected molecular weights for all EWI-F variants except for EWI-F_ΔIg1-4_ (Fig. 3F), suggesting that EWI-F_ΔIg1-4_ was possibly not expressed. Nevertheless, the co-purification of EWI-F_ΔIg1-5_ with CD9 indicates that the first five Ig-like domains of EWI-F are not essential for maintaining the CD9 - EWI-F interaction. This is in agreement with previous studies that identified the transmembrane helix of EWI-F as the main CD9 and CD81-interacting region (36, 38). Based on these co-purification experiments, we chose to perform our subsequent cryo-EM experiments with construct EWI-F_ΔIg1-5_, which harbors only a single Ig-like domain (Ig6) and thus presumably has the least conformational freedom of all tested EWI-F constructs.

### Cryo-EM of EWI-F_ΔIg1-5_ - CD9 - 4C8 complex

We next expressed Strep-tagged, full-length CD9 (without GFP) together with EWI-F_ΔIg1-5_ and purified the complex in digitonin in sufficient quantities for cryo-EM experiments. An EWI-F_ΔIg1-5_ dimer in complex with one or two CD9 molecules comprises <100 kDa of protein mass, making it a challenging sample for cryo-EM. We therefore investigated whether nanobody 4C8 could be employed to create a larger protein particle with more extra-membrane (or extra-micelle) features. To determine if nanobody 4C8 could still bind CD9 complexed to EWI-F_ΔIg1-5_, we incubated the complex with an excess of 4C8 and subjected it to a SEC-purification step. The resulting SEC-elution profile showed a monodisperse peak for the CD9 - EWI-F_ΔIg1-5_ complex (Fig. 4A); SDS-PAGE analysis of the peak then confirmed the presence of 4C8 (Fig. 4B), indicating that the epitope recognized by 4C8 is still accessible and that 4C8 does not disrupt the association between EWI-F_ΔIg1-5_ and CD9. The EWI-F_ΔIg1-5_ - CD9 - 4C8 complex was monodisperse in size as assessed by Trp-fluorescence assisted SEC (Fig. 4C), and thus suitable for analysis by single-particle cryo-EM. We collected a cryo-EM dataset and observed homogeneously sized-protein particles distributed in vitreous ice (Fig 4D), which allowed more accurate particle picking than for the previous dataset with full length EWI-F (Fig. 3D). 2D-class averages of the picked particles revealed expected density features, with protein domains protruding from a disk-shaped micelle region (Fig. 4E). Image processing in Relion yielded reconstructed density maps at a maximum global resolution of ~8.6 Å (Fig. 5A - C, Fig. S4, Table 2), although protein regions of the map exhibit a higher local resolution (Fig. S4B).

**Figure 4:**
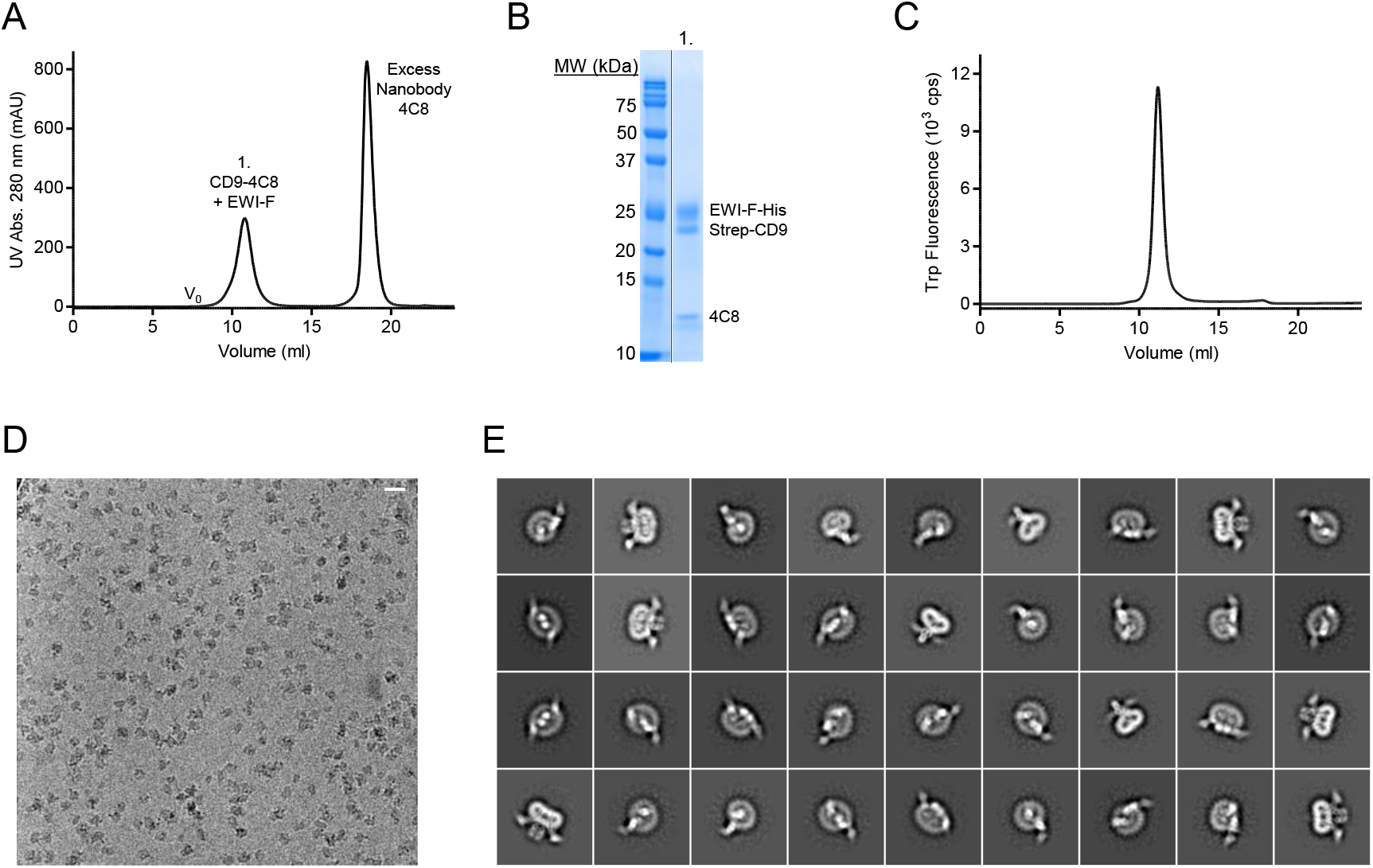
Cryo-EM sample preparation and imaging of EWI-F_ΔIg1-5_ - CD9 - 4C8. (A) Size-exclusion chromatography elution profile of co-expressed CD9 and EWI-F_ΔIg1-5_, after preincubation with a large excess of nanobody 4C8. (B) SDS page gel of peaks 1 of the SEC elution from panel A. (C) Analytical fluorescence-assisted SEC elution profile of the EWI-F_ΔIg1-5_-CD9-4C8 complex used for cryo-EM. (D) Cryo-EM micrograph depicting EWI-F_ΔIg1-5_-CD9-4C8 particles in vitreous ice. The scale bar length is 200 Å. (E) Selected 2D-class averages generated through Relion. The box size is 309×309 Å.

**Figure 5:**
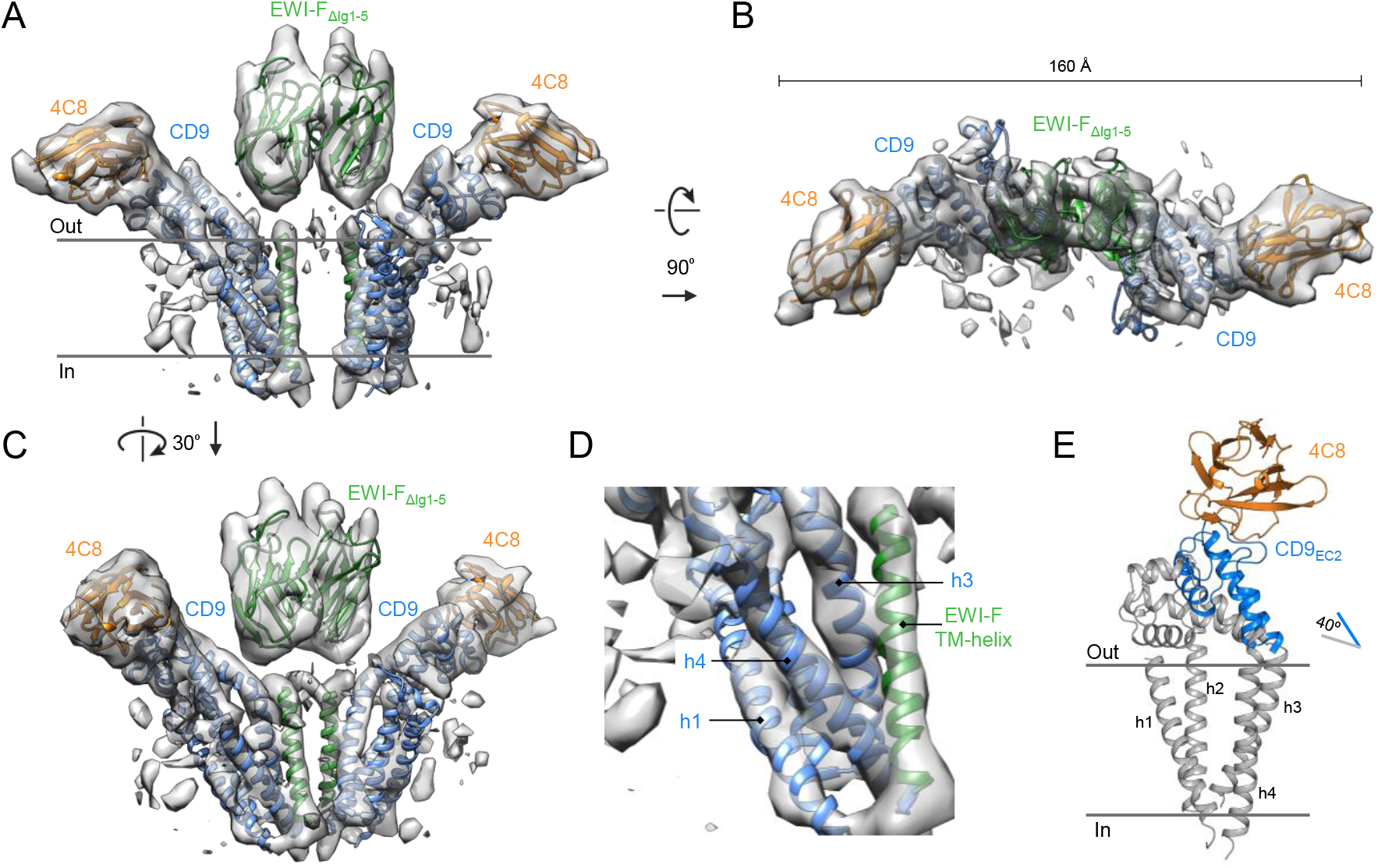
Cryo-EM structure of EWI-F_ΔIg1-5_ - CD9 - 4C8. (A - C) Sharpened, local-resolution filtered cryo-EM density map of EWI-F_ΔIg1-5_ - CD9 - 4C8 fitted with structures of CD9_EC2_ - 4C8, the TMD of CD9 (6K4J) and homology models of EWI-F, as viewed parallel to the membrane as a sideview (A), orthogonal to the membrane from the extracellular side (B), or as a sideview rotated by 30° (C). (D) Zoom of the major interaction region in a CD9 - EWI-F hetero-dimer. Membrane helices are annotated. (E) Overlay of the 4C8-bound CD9_EC2_ as oriented in the cryo-EM structure with the structure of CD81 (grey, pdb 5TCX).

**Table 2:**
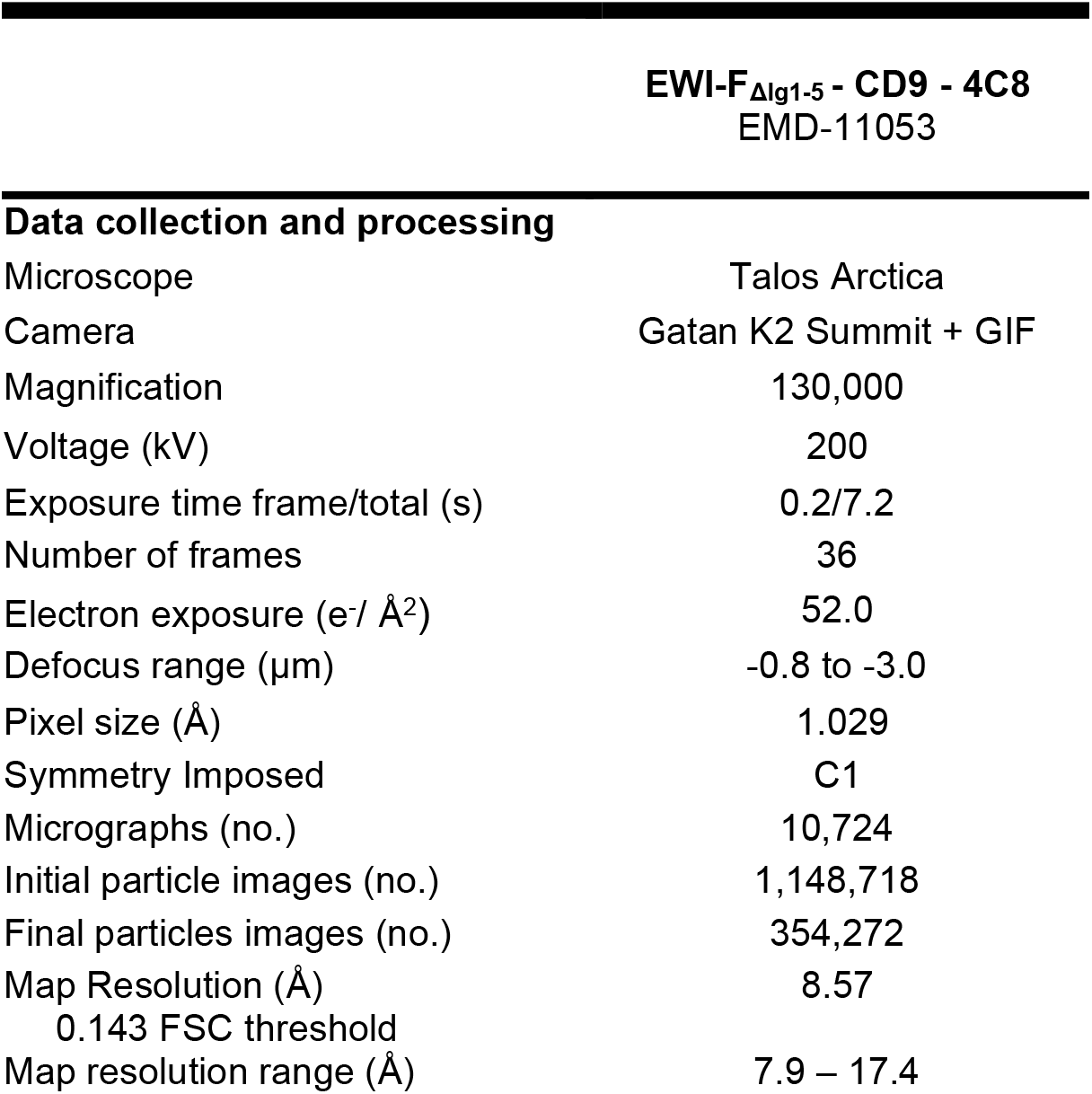
Cryo-EM data collection and processing.

### Complex architecture and flexibility

The ~8.6-Å resolution cryo-EM map revealed a hetero-tetrameric arrangement of CD9 - EWI-F_ΔIg1-5_, with a central EWI-F_ΔIg1-5_ dimer flanked by 4C8-bound CD9 on each side (Fig. 5A - C). Although the resolution of the reconstruction did not allow for *de novo* modelling, density regions corresponding to protein domains of 4C8-bound CD9 and EWI-F could be distinguished, and rigid-body fits of structures of CD9_EC2_ - 4C8 (Fig. 1A), the TMD of CD9 (pdb 6K4J) and homology models for the EWI-F Ig6 (chain b of pdb 1F3R) and TM-helix (pdb 5EH4) were in good agreement with the density map (Fig. 5A - C). Consistent with a TMD interface between CD9 and EWI-F_ΔIg1-5_ (36, 38), we observed that the EWI-F TM-helix resides in close proximity to CD9-helix h3 at the membrane center and to CD9-helix h4 at the cytoplasmic side of the membrane (Fig. 5D). The two TMD regions of the complex, each comprising the four helices of CD9 and the TM helix of EWI-F, are separated and do not interact, but instead are bridged by the extracellular dimeric Ig6 domain of the two EWI-F molecules. Overall, CD9 and EWI-F share no extensive interface through their extra-membrane domains, with no major contact areas between the EC2 domains of CD9 and EWI-F-Ig6. The 4C8-bound C and D loops of the EC2 orient away from EWI-F. Compared to the structure of CD81, the EC2 is rotated upwards by ~40°, resulting in an open conformation of CD9 when in complex with EWI-F (Fig. 5E).

Recently, a cryo-EM density map has been reported of CD9 in complex with an EWI-F homolog, EWI-2. Both the CD9 - EWI-2 map and our CD9 - EWI-F map were solved at moderate resolutions, limiting a detailed comparison of both reconstructions. However, both reconstructions reveal a similar hetero-tetrameric complex, with a central EWI-protein dimer and two CD9 molecules on each side (Fig. S5). Thus, CD9 interacts with its major partners EWI-F and EWI-2 in a comparable protein arrangement.

During the EM-image processing, it became apparent that the imaged EWI-F_ΔIg1-5_ - CD9 - 4C8 complex adopts a range of conformations. A typical 3D classification experiment with four major classes is shown in Fig. 6. The 4C8 - CD9 - EWI-F_ΔIg1-5_ - EWI-F_ΔIg1-5_ - CD9 - 4C8 composition suggests that the complex might arrange as a C2-symmetric particle. However, we observed no two-fold rotation axis in any of the four classes shown in Fig. 6, nor in any further sub-classifications. Class #1 resembles the orientation of the map presented in Fig. 5A and shows a pseudo two-fold arrangement, with a 9° deviation in anti-parallel orientation of the two 4C8-bound CD9 copies in the complex. In contrast, classes #2 – #4 show a much larger deviation from an anti-parallel CD9 configuration, i.e. a deviation of 49° is observed in class #4. A morph of 4C8-bound CD9 models fitted in the different classes reveals the putative conformational changes of CD9 in the complex (Video S1).

**Figure 6:**
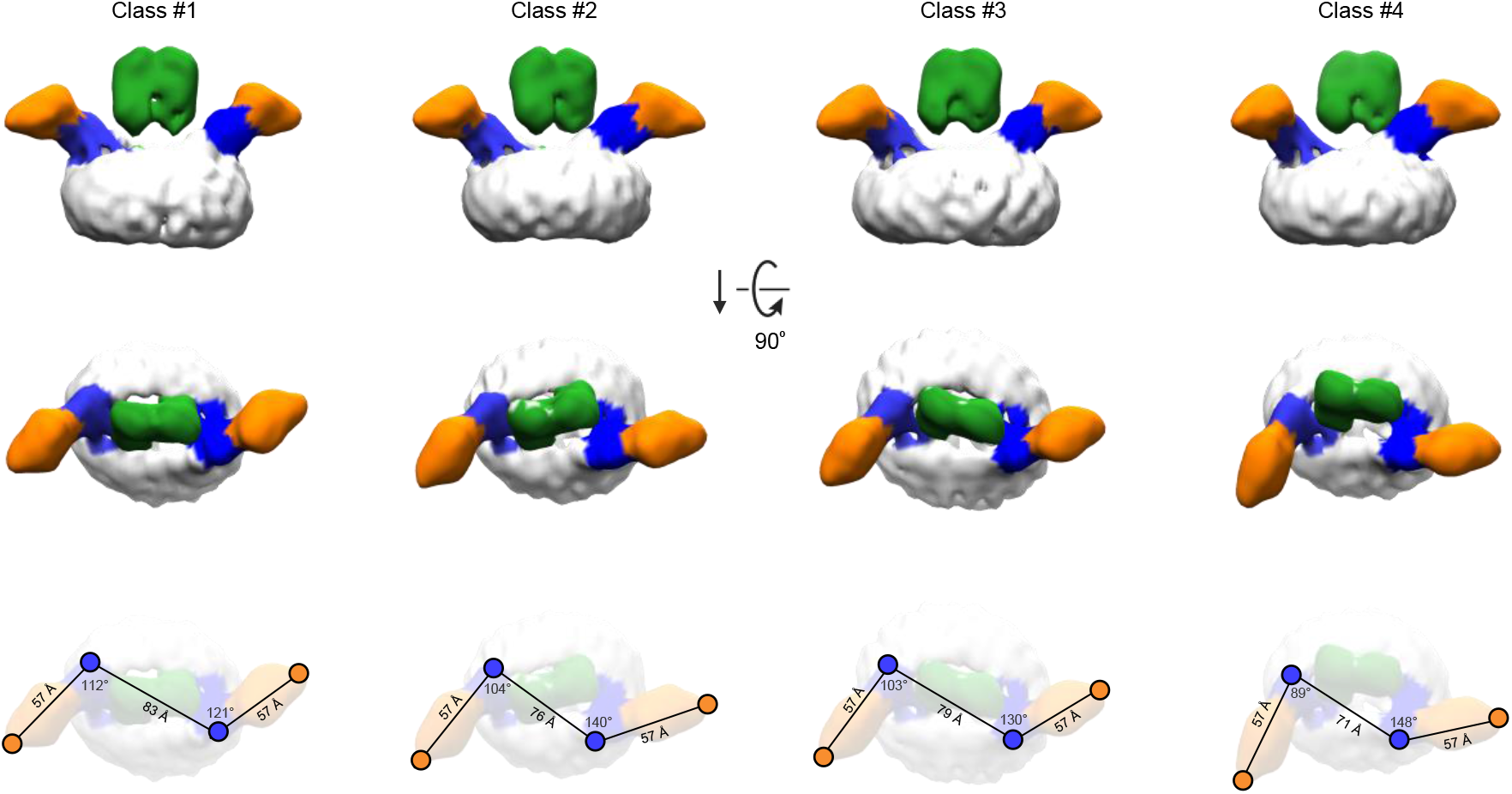
Flexibility of the EWI-F_ΔIg1-5_ - CD9 - 4C8 revealed by 3D classifications. Four classes obtained from a single 3D classification run are shown (see also Fig. S4). The density regions corresponding to the Ig6-domain of EWI-F, CD9_EC2_, nanobody 4C8 and the digitonin micelle are colored green, blue, orange and white, respectively. The top row depicts the four classes parallel to the membrane as a sideview. The middle row shows the classes orthogonal to the membrane from the extracellular side. The bottom row depicts the density maps in the same orientation as in the middle row, with annotated distances and angles between CD9_EC2_-residue K135 and 4C8-residue S121.

## Discussion

Tetraspanin-mediated interactions that drive the clustering of single-pass membrane receptors into TEMs in plasma membranes have remained obscure. Structural insights into tetraspanins have so far encompassed the large extracellular EC2 loop of CD81 (23, 40–43), full-length CD81 (24) and, more recently, full-length CD9 and a low-resolution cryo-EM map of CD9 in complex with EWI-2 (25).

The three crystal structures, presented here, reveal an overall architecture of CD9_EC2_ similar to that of the homologous CD81_EC2_, except for a more extended arrangement of the C-loop and, in particular, D-loop regions. In structures of CD81, these regions adopt more helical conformations, though partially unfolded arrangements are also observed in antibody-bound CD81_EC2_ structures (43). The D-loop conformations in CD9_EC2_ range from a partial helical arrangement in the nanobody-bound structures to an extended loop with no secondary structure elements in the domain-swapped CD9_EC2_ structure without nanobody (Fig. 2B). The homo-dimerizing D-loops in between two domain-swapped CD9_EC2_ dimers (Fig. 2A) could explain the proposed homo-dimerization interface in the homologous CD81, as mediated by D-loop residues P176 – F186 (44–46) (which correspond to P168 – T177 in CD9). Overall, the ability of the D-loop, which is sequence variable among tetraspanins, to adopt multiple conformations is preserved in the structures of CD9_EC2_ and CD81_EC2_. This observation supports the hypothesis that the EC2 head region can be tailored to facilitate interactions with specific partner proteins (49). In the full-length crystal structure of CD81, molecules pack in an anti-parallel fashion through hydrophobic interaction of their TM regions, resulting in a non-physiological up-down arrangement (24). Although the molecules in the crystal structure of full-length CD9 (25) align in a more physiologically relevant manner, potential homo-dimerizing contacts between EC2 fragments were abrogated due to the deletion of D-loop residues T175 – K179 (50). When taken together, the variable D-loop conformations observed in crystal structures are consistent with the representation of the D-loop as an ‘interaction hub’ for homo- and hetero-oligomerization in the head region of tetraspanins, as proposed earlier (5).

The cryo-EM map of CD9 in complex with EWI-F_ΔIg1-5_ and nanobody 4C8 revealed a core CD9 – EWI-F - EWI-F - CD9 hetero-tetramer arranged in a linear fashion. In this complex, nanobodies 4C8 bind to the CD9 on either side without contacting the EWI-F molecules. Both CD9_EC2_ segments are tilted by ~40° with respect to their TMD region (Fig. 5E), yielding an open conformation that is comparable to those predicted by MD simulation of CD81 and CD9 starting from the closed conformations observed in the crystal structures (24, 25). CD9 predominantly contacts EWI-F through its TMD region, where CD9 helices h3 and h4 interact with the single-pass TM-helix of EWI-F (Fig. 5D), which is in agreement with prior biochemical data that identified membrane helix h4 as a critical interface for binding EWI-F (36, 38). The TMD regions of both CD9 - EWI-F hetero-dimers are separated by more than 10-Å distance and make no or minimal direct contacts. Instead, dimerization of the two hetero-dimers into a tetramer is dominated by contacts between the extracellular Ig6 domains of EWI-F_ΔIg1-5_. Consistent with the absence of an intra-membrane interface between the two CD9 - EWI-F heterodimers and no major extra-membrane contacts between CD9 and EWI-F (Fig. 5), the overall arrangement is highly flexible and we observed a range of linear-arranged tetrameric complexes, ranging from an extended conformation to one that is bent by ~50° (Fig. 6, Video S1).

Most recently, Umeda et al. reported a cryo-EM map of CD9 in complex with EWI-2 (25). Overall, the structural arrangements of CD9 - EWI-F_ΔIg1-5_ and CD9 - EWI-2 complexes are similar (Fig. S5), in line with the partially overlapping cellular functions of EWI-F and EWI-2 (31, 37). As with CD9 - EWI-F, major contacts are observed between h3 and h4 of CD9 and the single-pass TM helix of EWI-2. However, the membrane helices of EWI-2 are in closer proximity to each other than those of EWI-F, which could be correlated with the presence of two more putative palmitoylation sites in EWI-F than EWI-2. Furthermore, for CD9 - EWI-2, only one conformation was reported, whereas the distinct shape of the CD9_EC2_ - 4C8 density enabled us to distinguish a range of straight to bent conformations for CD9 - EWI-F_ΔIg1-5_ (Fig. 6, Video S1). However, given the limited resolution of the cryo-EM maps of both cases, flexibility appears to be an inherent feature of the two complexes and a similar bending of CD9 - EWI-2 cannot be excluded.

CD9, CD81 and other tetraspanins interact with various single-pass TM proteins that are known to homodimerize through their extra-membrane domains. Examples of homodimerizing partners of tetraspanins (besides EWI-F and EWI-2) include: DPP4 and ADAM17 for CD9; EGFR for CD82 (51); and hetero-dimeric integrins for CD151 (52–54). The observed linear-arranged tetramers of CD9 – EWI-F and CD9 – EWI-2 indicate that the interactions that govern complex formation are sequentially hydrophobic-hydrophilic-hydrophobic, with tetraspanin partner-protein interactions mediated by the TMDs on either side and protein-partner homo-dimerization via their ectodomains in the center. Previous studies mapped a tetraspanin homo-dimerization site (for CD9) to membrane helices h1 and h2 (55) and, for CD81, to the variable D-loop in the EC2 head domain. Both helices h1 and h2 and the D-loops are positioned on either end of the linear-arranged tetramers and orient outwards posed for putative interactions (Fig. 5A). Thus, the observed hetero-tetrameric arrangements (Fig. 6) suggest a simple ‘concatenation model’ for forming transient higher-order assemblies of end-to-end attached tetramers yielding small linear or circular structure (Fig. 7), which may explain the occurrence of TEMs and their highly dynamic nature. This single-particle cryo-EM-derived model is in agreement with scanning-EM data on immunogold-labeled CD81 and CD9 expressed on cells, which showed both tightly packed clusters and linear tetraspanin assemblies (56, 57). Thus, the concatenation model provides a structural basis for further studying the formation and signaling of TEMs in the plasma membrane.

**Figure 7:**
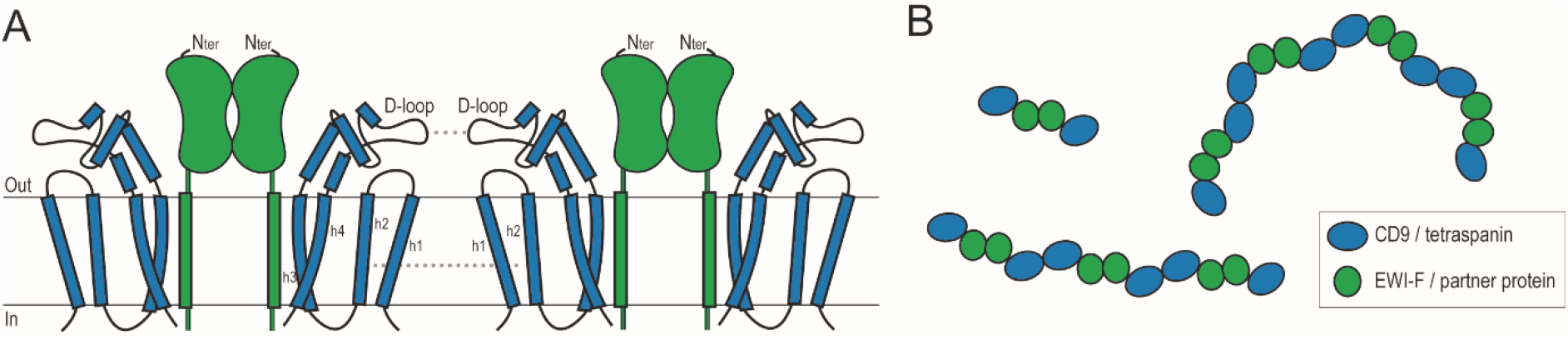
Concatenation model of TEM formation. (A) Cartoon model for the formation of higher-order oligomers based on the hetero-tetrameric CD9 - EWI-F arrangement observed in the cryo-EM data and biochemical interaction studies. (B) Top-view model, shown orthogonal to the membrane, of putative linear and circular TEM assemblies based the straight and bent conformations adopted by the CD9 - EWI-F complex and the oligomerization model in panel A.

## Experimental procedures

### Chemicals

All chemicals were purchased from Sigma-Aldrich unless specified otherwise.

### Construct design

The cDNA encoding for CD9 and EWI-F was obtained as described previously (39). Briefly, codon-optimized cDNA for mammalian cell expression, encoding for human CD9 was purchased from Geneart. Full-length CD9 was cloned in a pUPE expression vector (U-Protein Express BV) with either a N-terminal Strep-II tag or a N-terminal Strep3-GFP tag with a TEV-protease site. The CD9-W6 construct (CD9-C9W,C78W,C79W,C87W,C218W,C219W), with all palmitoylated cysteines mutated to tryptophan, was cloned as described (39). The construct encoding for the EC2 domain of CD9 (CD9_EC2_, residues K114 – N191) was generated with a PCR reaction using the Q5-PCR kit (NEB). The CD9_EC2_ construct was cloned in a pUPE expression vector with a N-terminal cystatin signal peptide and a C-terminal 6x-His tag. cDNA encoding for full-length human EWI-F was obtained from Source BioScience. EWI-F truncations were generated using PCR. Construct boundaries of EWI-F_ΔIg1_ (H146 – D879), EWI-F_ΔIg1-2_ (Q275 – D879), EWI-F_ΔIg1-3_ (E405 – D879), EWI-F_ΔIg1-4_ (N540 – D879) and EWI-F_ΔIg1-5_ (T685 – D879) were based on Uniprot domain assignments (UniprotKB - Q9P2B2). All EWI-F constructs were cloned in a pUPE expression vector with a N-terminal cystatin signal peptide and a C-terminal 6x-His tag.

### Nanobody selection and production

Phage library construction and anti-CD9 nanobody selections were carried out as described before and resulted in the selection of two nanobodies: 4C8 and 4E8 (39). DNA sequences of clones 4C8 and 4E8 were ligated into a modified pHEN6 vector with a C-terminal thrombin-cleavable 6xHis tag and an N-terminal pelB leader sequence for periplasmic secretion (gift from dr. F. Opazo, University of Göttingen Medical Center, Göttingen). Large scale nanobody productions were performed overnight at 25°C using an Bioflo 115 Bioreactor (New Brunswick) as previously described (37). Bacterial biomass was harvested by centrifugation at 5,400*g* and His-tagged nanobodies were purified using HisTrap™ column chromatography (GE Healthcare) followed by a buffer exchange step to PBS using a HiTrap™ desalting column (GE Healthcare). Nanobodies were stored in PBS at −20 °C until further use. Nanobody 4C8 was treated with Thrombin (0.5U/100μg; GE Healthcare) for 16 hrs at 22°C, to remove the 6xHis purification tag. 4C8 was then injected into a Superdex75 10/300 column (GE Healthcare) pre-equilibrated in buffer A (25 mM HEPES pH 7.5, 150 mM NaCl), and monodisperse fractions were collected. The tag of nanobody 4E8 was not removed prior to crystallization.

### Determination of nanobody binding affinity

The binding affinity of 4C8 and 4E8 on CD9 was determined both on purified CD9 and on CD9 endogenously expressed on HeLa cells as described in detail (39). Binding was carried on either 100 ng wtCD9-3Strep captured on Strep-Tactin® coated wells; or HeLa cells seeded in 96-well plates. On the day of the assay, target was incubated with serial dilutions of nanobodies for 2 hours either in 2% (w/v) Bovine Serum Albumin (BSA) in buffer A supplemented with 0.025% (w/v) N-Dodecyl-ß-D-maltoside (DDM) at rt in the case of purified protein; or in Binding buffer (1% w/v BSA and 25 mM HEPES pH 7.2 in DMEM without phenol red; Lonza Netherlands B.V.) at 4°C in the case of HeLa cells, in triplicate. Nanobodies were detected after incubation with rabbit anti-VHH antibody (1:1000 in 2% w/v BSA in PBS; clone k1216, QVQ B.V.) for 1 hour at room temperature, followed by 1 hour incubation with goat anti-Rabbit IRDye800CW (1:1000 in 2% w/v BSA in PBS; LI-COR Biosciences) at rt.

For cell binding assays, cells were fixed with 4% formaldehyde prior to the incubation with the detecting antibodies. Finally, IRDye800CW fluorescent signal at 800 nm was detected with Odyssey® Infrared Imager (LI-COR Biosciences). The measured fluorescent intensities were normalized, setting as 100% the mean of intensities detected at the higher nanobody concentrations, and plotted (mean ± SEM) over protein concentration using GraphPad Prism 7. A non-linear regression curve for one-site specific binding was fitted in order to determine apparent binding affinity (K_D_) of the different nanobodies.

### CD9_EC2_ expression and purification

CD9_EC2_ was recombinantly expressed in HEK293-EBNA cells (provided by U-Protein Express BV). Cells were grown at 37°C and harvested after six days. The cells were centrifuged at 1,000*g* for ten minutes, after which the medium was collected. Two different purification strategies were then employed:

1. For the protein sample used for the crystallization of CD9_EC2_ in the absence and presence of nanobody 4C8, the medium was loaded overnight onto a ~5 ml anti-CD9 AR40A746.2.3 affinity column (7.5 mg antibody/ml CNBr Activated Sepharose 4B beads; GE Healthcare) in line of an AKTA explorer (GE). After loading, unbound proteins were washed away and bound CD9_EC2_ was eluted using 0.2 M Glycine, 150 mM NaCl pH 2.5 and neutralized with 1:10 volume of 1M Tris pH 9.0. A size-exclusion step on a Superdex 200 10/300 Increase column equilibrated in buffer A (25 mM HEPES pH 7.5, 150 mM NaCl) was performed. Next, another affinity-purification step was performed on a ~5 ml anti-CD9 AT1412dm (58) affinity column (7.5 mg antibody/ml CNBr Activated Sepharose 4B beads). After loading, unbound proteins were washed away and bound CD9_EC2_ was eluted with 0.2 M Glycine, 150 mM NaCl pH 2.5 and neutralized 1:10 volume of 1M Tris pH 9.0. A final size-exclusion step on a Superdex 200 10/300 Increase column equilibrated in Buffer A was performed prior to crystallization.
2. For the protein sample used for the crystallization of CD9_EC2_ in complex with nanobody 4E8, the cell medium was incubated with Ni-Sepharose Excel beads (GE Healthcare) at 4°C for 2 hours. The beads were washed for 10 column volumes in buffer B (50 mM Tris pH 7.8, 500 mM NaCl) with 10 mM imidazole and CD9_EC2_ was subsequently eluted from the beads with 300 mM imidazole in buffer B. The protein sample was then concentrated and incubated with 2 mM DTT (final concentration) for ~16 hours to remove intermolecular, non-native disulfide bonds. CD9_EC2_ was then injected on a Superdex75 10/300 column (GE Healthcare) pre-equilibrated in buffer C (20 mM Tris pH 8, 150 mM NaCl). Monomeric protein fractions were pooled based on non-reducing SDS-PAGE. To form a complex with nanobody 4E8, 230 μl of CD9_EC2_ at 5 mg/ml was mixed with 250 μl of 4E8 (7 mg/ml, in PBS) and incubated for 30 minutes. The mixture was injected on a Superdex75 10/300 column pre-equilibrated in buffer C and fractions containing both CD9_EC2_ and 4E8 were collected and concentrated to ~8.0 mg/ml.

### Crystallization and data collection

CD9_EC2_ (10 mg/ml) was crystallized in sitting drop in 33% w/v pentaerythritol propoxylate, 0.2 M KCl, 0.1 M sodium citrate at pH 6.0. No further cryoprotectant was used.

For the CD9_EC2_-4C8 complex, CD9_EC2_ (12 mg/ml in buffer A) was mixed with 4C8 (5.9 mg/ml in buffer A) in a 1:1 molar ratio. Crystals grew in sitting drop in 0.095 M sodium citrate pH 5.6, 5% (v/v) glycerol, 19% (v/v) isopropanol, 20% (w/v) PEG 4,000. Crystals were cryoprotected by soaking in reservoir solution supplemented with 25% glycerol (final concentration).

The CD9_EC2_ - 4E8 complex was purified as described above and concentrated to 8.0 mg/ml. Crystals grew in hanging drop in 0.2 M sodium acetate, 0.1 M Tris pH 8.0, 30% (w/v) PEG 4,000 and were cryoprotected by soaking in reservoir solution supplemented with 20% (v/v) ethylene glycol.

For the crystallization of nanobody 4C8 alone (5.1 mg/ml), crystals grew in sitting drop in 0.1 M HEPES pH 7.5, 20% (w/v) PEG 8,000 and were cryoprotected by soaking in reservoir solution supplemented with 30% (v/v) PEG 400.

All crystals were flash frozen in liquid nitrogen immediately after harvesting. Diffraction data were collected at Diamond Light Source (DLS) on beamlines I-03 (CD9_EC2_ - 4C8), I-04 (CD9_EC2_) and I-04-1 (CD9_EC2_ - 4E8), or at the European Synchrotron Radiation Facility (ESRF) on beamline ID23-2 (4C8 alone).

### CD9_EC2_ crystal data processing and refinement

The triclinic CD9_EC2_ crystal was twinned with a twofold rotation about ***a\****+***b\**** as the twinning operation. Consequently, two orientation matrices were used for integration with the Eval15 software (59). In the prediction of reflection profiles an isotropic mosaicity of 1.1° and a mica expansion of 0.077 along ***a\****+***b\**** was assumed. The resulting reflection file contained 21.5% overlapping reflections belonging to both twin domains. Initial de-twinning was performed with the TWINABS software (60). These data were used for structure solution by molecular replacement, using PHASER (61) within the CCP4 software suite (62), using the model of CD9_EC2_ from the previously solved AT1412dm Fab-CD9_EC2_ complex (58) as model. The structure was iteratively refined using Refmac5 (63) alternated with model improvement in COOT (64). Local non-crystallographic symmetry (NCS) restraints were maintained during refinement. The calculated structure factors from this refinement were then used for the final de-twinning of the overlapping data. The scale factor of 6.2147 * exp(−8.19544*sin(θ/λ)^2^) between the two twin domains was determined with XPREP (65), based on the non-overlapping reflections. Interframe scaling of the de-twinned data was performed with SADABS and merging was performed with the CCP4 suite. Final refinement rounds in Refmac5 using the latest data yielded R_work_/R_free_ = 23.9/27.9% and the structure was deposited in the RCSB Protein Data Bank under accession code 6RLR.

### CD9_EC2_ - 4C8 crystal data processing and refinement

Diffraction images were processed using DIALS (66) and the integrated reflection data were truncated anisotropically using the STARANISO web server (67). The structure was solved by molecular replacement using PHASER with the CD9_EC2_ structure and a nanobody homology model obtained through the SWISS-MODEL server (68) as search models. The structure was iteratively refined using Refmac5 or Phenix (69) alternated with manual model improvement in COOT. The final refinement in Phenix yielded R_work_/R_free_ = 20.6/24.8% and the structure was deposited in the RCSB Protein Data Bank under accession code 6Z20.

### 4C8 crystal data processing and refinement

Diffraction images were processed using Eval15 (59). The structure was solved by molecular replacement using PHASER with the nanobody in chain B of PDB 5IMK as the search model. The nanobody residues were manually adjusted to the 4C8 amino-acid sequence in Coot. The structure was then iteratively refined using Refmac5 or Phenix, alternated with manual model improvement in COOT. The final refinement in Phenix yielded R_work_/R_free_ = 20.1/23.1% and the structure was deposited in the RCSB Protein Data Bank under accession code 6Z1Z.

### CD9_EC2_ - 4E8 crystal data processing and refinement

For the CD9_EC2_ - 4E8 dataset, the autoprocessed and anisotropical-truncated (autoproc-staraniso) reflection data file provided by DLS was used. The structure was solved by molecular replacement using PHASER with the CD9_EC2_ - 4C8 structure as search model. The 4C8 residues were replaced with the corresponding 4E8 residues and the CDR regions of the nanobody were manually built in Coot. The structure was then iteratively refined using Refmac5 or Phenix, alternated with model improvement in COOT. The final refinement in Phenix yielded R_work_/R_free_ = 15.1/19.0% and the structure was deposited in the RCSB Protein Data Bank under accession code 6Z1V.

### Large-scale expression and purification of CD9 and full-length EWI-F

N-Strep3-GFP-tagged CD9-W6 and full-length EWI-F were transiently expressed in 2L Epstein-Barr virus nuclear antigen I (EBNA1)-expressing HEK293 cell cultures (HEK293-EBNA, provided by U-Protein Express BV). Cells were grown at 37°C and harvested after four days. All subsequent steps were carried out at 4°C. Cells were washed in PBS and then lysed in buffer containing 50 mM Tris pH 7.8, 150 mM NaCl, 1% (w/v) N-dodecyl-ß-D-maltoside (DDM, Anatrace) and protease inhibitor cocktail (Roche) for two hours. The lysed sample was ultracentrifuged for 45 min at 100,000 *g* to remove insoluble membranes and cell debris. The supernatant was incubated with Streptactin resin (GE Healthcare) for two hours and the resin was washed with 20 column volumes of buffer D (50 mM Tris pH 7.8, 150 mM NaCl, 0.025 % (w/v) DDM). Protein was eluted from the resin with buffer D supplemented with 3.5 mM desthiobiotin. The eluted fractions were pooled, concentrated in a 100 kDa concentration device (Amicon) and injected on a Superose6 10/300increase column (GE Healthcare) pre-equilibrated in buffer D. Fractions containing both CD9 and EWI-F were pooled and concentrated to ~3.3 mg/ml.

### Small-scale expression and purification of CD9 with EWI-F variants

N-Strep3-GFP-tagged CD9-W6 was co-transfected with EWI-F, EWI-F_ΔIg1_, EWI-F_ΔIg1-2_, EWI-F_ΔIg1-3_, EWI-F_ΔIg1-4_ or EWI-F_ΔIg1-5_ in 25 ml HEK293-EBNA cell cultures (provided by U-Protein Express BV). Cells were grown at 37°C and harvested after four days. Cells were washed in PBS and then lysed in buffer containing 50 mM Tris pH 7.8, 150 mM NaCl, 1% (w/v) DDM, 0.5% (w/v) digitonin (Calbiochem) and protease inhibitor cocktail (Roche) for two hours. The supernatant was incubated with Streptactin resin (GE Healthcare) for two hours and the resin was washed with buffer E (50 mM Tris pH 7.8, 150 mM NaCl, 0.08 % (w/v) digitonin) in spin columns. Protein was eluted from the resin with buffer E supplemented with 3.5 mM desthiobiotin. Complex formation between CD9 and the co-transfected EWI-F variants was assessed by SDS PAGE.

### Large-scale expression and purification of CD9 and EWI-F_ΔIg1-5_ with nanobody 4C8

N-StrepII-tagged, wild type CD9 and EWI-F_ΔIg1-5_ were expressed in 3L HEK293-EBNA GNT1-cell cultures. Cells were grown at 37°C and harvested after four days. All subsequent steps were carried out at 4°C. Cells were washed in PBS and then lysed in buffer containing 50 mM Tris pH 7.8, 150 mM NaCl, 1% (w/v) digitonin 0.2% (w/v) DDM and protease inhibitor cocktail (Roche) for two hours. The lysed sample was ultracentrifuged for 45 min at 100,000 *g* to remove insoluble membranes and cell debris. The supernatant was incubated with Streptactin resin (GE Healthcare) for two hours and the resin was washed with 20 column volumes of buffer E (50 mM Tris pH 7.8, 150 mM NaCl, 0.08 % (w/v) digitonin). Protein was eluted from the resin with buffer E supplemented with 3.5 mM desthiobiotin. The eluted fractions were pooled, concentrated in a 100 kDa concentration device (Amicon) to ~1.5 mg/ml and incubated with EndoHF (NEB) at a volume to volume ratio of 1:20 for 2 hours to remove the N-linked glycan of EWI-F_ΔIg1-5_. The complex was then incubated with a large excess of nanobody 4C8 (in PBS buffer with 0.08% (w/v) digitonin) and injected on a Superdex200 10/300increase column (GE healthcare) pre-equilibrated in buffer E. Fractions containing the EWI-F_ΔIg1-5_ - CD9 - 4C8 complex were pooled and concentrated to ~3 mg/ml.

### Cryo-EM grid preparation and data collection

2.8 μl of CD9 - EWI-F_Full-Length_ (3.3 mg/ml) or EWI-F_ΔIg1-5_ - CD9 - 4C8 (3 mg/ml) was pipetted onto a glow-discharged R1.2/1.3 200 mesh Au holey carbon grid and then plunge-frozen in a liquid ethane/propane mixture using a Vitrobot Mark IV (Thermo Fisher Scientific). The blotting was performed at 20 °C for four seconds with blot force 0 (for CD9 - EWI-F_Full-length_) or blot force 1 (for EWI-F_ΔIg1-5_ - CD9 - 4C8).

All data was collected on a 200 kV Talos Arctica microscope (Thermo Fisher Scientific) equipped with a K2-summit detector (Gatan) and a post-column 20 keV energy filter, using EPU automated-data collection software (Thermo Fisher Scientific). Movies for the CD9-EWI-FFull-Length dataset were collected in super-resolution mode (binned pixel size 1.03 Å) in 28 frames for 7 seconds, with an electron exposure of 1.82 e^−^/Å^2^/frame (total exposure 50.9 e^−^/Å^2^). Movies for the EWI-F_ΔIg1-5_-CD9-4C8 dataset were collected in counting mode (pixel size 1.03 Å) in 36 frames for 7.2 seconds, with an electron exposure of 1.45 e^−^/Å^2^/frame (total exposure 52.0 e^−^/Å^2^).

### Cryo-EM image processing

For the CD9 - EWI-F_Full-Length_ dataset, 379 micrographs were imported in the Relion3.0beta pipeline (70). The super-resolution mode recorded micrographs were binned 2x, gain corrected and motion corrected using MotionCor2 (71), and GCTF (72) was used to estimate the contrast transfer function for each micrograph. 880 particles were manually picked and 2D classified. The resulting 2D-class averages were then used as templates for automated particle picking in Relion (73), yielding 29,216 particles, that were binned 2x upon particle extraction. These particles were subjected to numerous 2D classification runs. Further processing was not attempted due to strong heterogeneity in the particles.

For the EWI-F_ΔIg1-5_ - CD9 - 4C8 dataset, 10,724 micrographs were imported in the Relion3.1beta pipeline. The micrographs were motion-corrected and gain-corrected using MotionCor2, and GCTF was used to estimate the contrast transfer function for each micrograph. 428 micrographs were discarded based on poor CTF estimations, yielding a total of 10,296 micrographs for further processing. The motion-corrected micrographs were imported in EMAN2 (74), and several hundred good particles and bad particles, as well as background images, were picked manually for training of the EMAN2 NeuralNet particle picker, which was subsequently used for automated particle picking. The obtained particle coordinates were imported back into the Relion pipeline and 1,148,718 particles were extracted and binned 3x (resulting pixel size 3.09 Å). The particles were 3D classified into four classes, and the 107,830 particles belonging to one obvious junk class were discarded. 1,040,888 particles were then subjected to two rounds of 2D classification into 200 classes, through which 352,979 particles were removed. The remaining 687,909 particles were 3D classified into 20 classes. The highest-populated class, comprising 354,272 particles, was 3D-auto refined using a mask and solvent flattening fourier shell correlations (FSCs), which yielded a map at a global resolution of 8.6 Å based on the gold-standard FSC = 0.143 criterion (75). This map was sharpened with a B-factor of −1200 Å^2^ and filtered based on local-resolution in Relion. Another 3D classification with the 687,909 particles (following the 2D classifications) was performed into five classes, while ignoring the CTFs until the first peak, meaning that CTF-amplitude correction was only performed from the first peak of each CTF onward. This strategy resulted in better particle separation in distinct classes. However, further sub-classifications, either with or without regions of the protein complex masked out, yielded density maps of worse quality and resolution. This suggested a continuous disorder in the protein complex rather than that few discrete conformations were adopted by EWI-F_ΔIg1-5_ - CD9 - 4C8. The particles were not un-binned as the Nyquist frequency (6.2 Å) was not reached in any refinement.

### Modeling in cryo-EM maps

Protein models were rigid-body fitted into the sharpened 8.6-Å local-resolution filtered EWI-F_ΔIg1-5_ - CD9 - 4C8 map, as well as in non-sharpened maps obtained through 3D classifications, using the ‘Fit in Map’ option in UCSF Chimera (76).

### Figure preparation

The crystal-structure figures were prepared using Pymol (Schrödinger). All figures containing cryo-EM density maps were generated using UCSF Chimera. The morph between four observed conformations of the EWI-F_ΔIg1-5_ - CD9 - 4C8 complex (Video S1) was made in UCSF Chimera and edited using Adobe Premiere Project. The cartoon models (Fig. 7) were prepared in Adobe Illustrator.

## Data availability

Data supporting the findings of this manuscript are available from the corresponding author (P.G.) upon reasonable request. The relevant cryo-EM density maps of the EWI-F_ΔIg1-5_ - CD9 - 4C8 dataset have been deposited in the Electron Microscopy Data Bank under accession number EMDB-11053. This deposition comprises the sharpened, local-resolution filtered map, unfiltered-half maps and four 3D-class averages. Model coordinates of the crystal structures have been deposited in the Protein Data Bank under accession numbers 6RLR (CD9_EC2_), 6Z20 (CD9_EC2_ - 4C8), 6Z1V (CD9_EC2_ - 4E8) and 6Z1Z (4C8).

## Author contributions

W.O, V.N. and P.G conceived the project and interpreted all data; K.T.X. and S.D. performed nanobody selection, purification and characterization; W.P., W.O. and J.M. purified soluble CD9_EC2_ protein; W.O., V.N. and J.M. performed crystallization experiments; W.O., V.N. and M.L. collected and processed crystal data; L.K-B and M.L processed and analyzed the non-merohedrally twinned crystallographic data; W.O., V.N. and N.M.P. built the protein models; W.O. purified all membrane proteins and performed the cryo-EM data collection and processing; P.M.P.B.H and P.G. acquired funding and provided project supervision; W.O. and P.G. wrote the manuscript with critical input from all authors.

## Acknowledgement

We thank the beamline scientists of Diamond Light Source (DLS) and the European Synchrotron Radiation Facility (ESRF) for assistance during data collection; we acknowledge W. Hemrika (U-Protein Express BV) for mammalian cell cultures; we thank S.C. Howes, C.T.W.M. Schneijdenberg and J.D. Meeldijk of the Utrecht EM-square for electron-microscope assistance and maintenance; we thank D. El Mazouni for fruitful discussions; we are grateful to A.B. van Spriel and S. van Deventer of Radboud UMC Nijmegen for proofreading of the manuscript. This work has been supported by the Netherlands Organization for Scientific Research (NWO), Fund NCI Technology Area (project no. 731.015.201) to P.M.P.B.H. and P.G.; and the Institute of Chemical Immunology (project 024.002.009) to P.G.

