## Supporting Note 1, Supporting Figures S1 - S5 for "Implications for tetraspanin-enriched microdomain assembly based on structures of CD9 with EWI-F"

[1]Crystal and Structural Chemistry, Bijvoet Centre for Biomolecular Research, Department of Chemistry, Faculty of Science, Utrecht University, The Netherlands.
[2]Cell Biology, Neurobiology and Biophysics, Department of Biology, Faculty of Science, Utrecht University, The Netherlands
[3]uniQure Biopharma, Amsterdam, The Netherlands

### Supporting information

#### Supporting Note 1

#### Figures S1 – S5

#### Supporting References 1 – 4

### Supporting note 1

**Diffuse scattering from $CD9_{EC2}$ crystals.**

Reciprocal space reconstructions were obtained from the diffraction images, using *IMG2HKL*, which is part of the EVAL package (1). Figure S3A shows three sections through reciprocal space. The crystal is twinned by a two-fold rotation along ***a*+b****. Figure S3B shows two twin domains with the twinning interface in the middle: a layer with base vectors c and a-b. Each layer is a possible twinning interface and diffuse streaks in the ***a*+b**** direction imply that the layers can also stack randomly. Starting from the middle layer, every fourth layer, belonging to the two twin structures, exactly overlap. Therefore, the structure can be indexed on a so-called stacking lattice (2, 3) with dimension 1/4c. On this lattice the twinned structure is completely ordered, causing reciprocal space reconstruction slices at l=4n to be ordered and all slices in between to have streaks in the direction ***a*+b****, i.e. the direction of packing disorder (Fig. S3B).

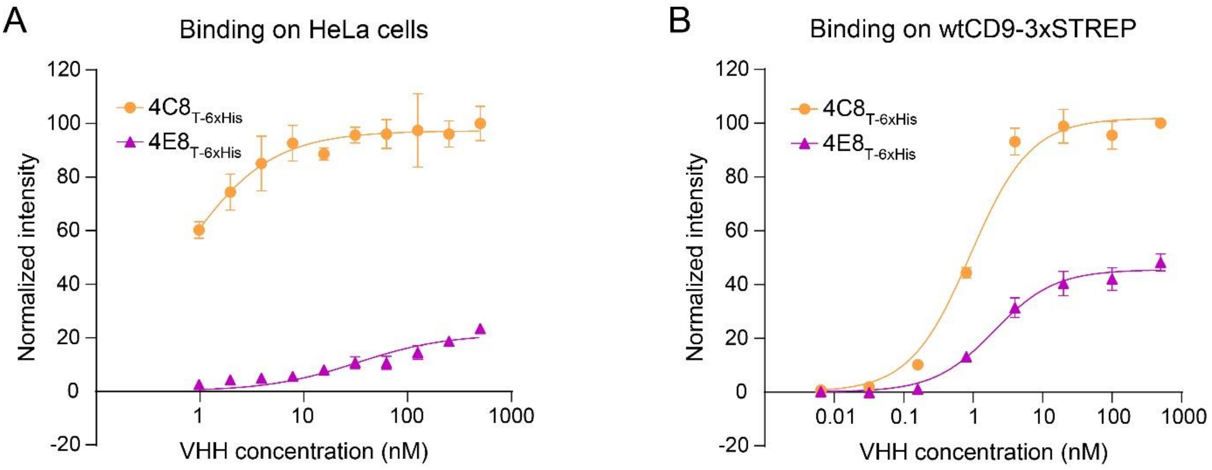


**Figure S1: Characterization of anti-CD9 nanobodies 4C8 and 4E8.** (A) Concentration-dependent binding curves of 4C8 and 4E8 binding to endogenous CD9 on HeLa cells. 4C8 and 4E8 exhibit affinities of 0.6 nM and 33 nM for CD9 on HeLa cells, respectively. (B) Concentration-dependent binding curves of 4C8 and 4E8 binding to detergent-purified, full-length CD9. 4C8 and 4E8 exhibit affinities of 0.9 nM and 2.0 nM for purified CD9, respectively. Comparable binding curves for 4C8 were described previously (4).

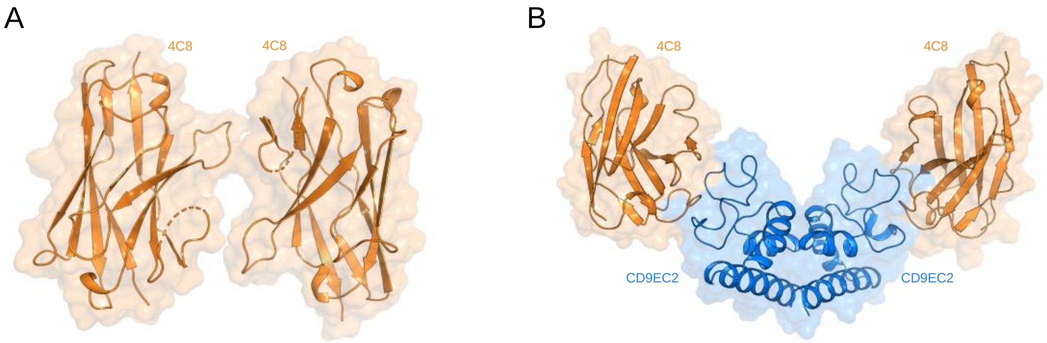


**Figure S2: Structures of 4C8 alone and bound to $CD9_{EC2}$.** (A, B) Asymmetric units of the of the structure of 4C8 alone (A) and the $CD9_{EC2}$ - 4C8 structure (B). 4C8 alone crystallized in space group $P2_1$ with two copies in the asymmetric unit. Residues 102 – 105 of CDR3 are not modelled in the 4C8 structure.

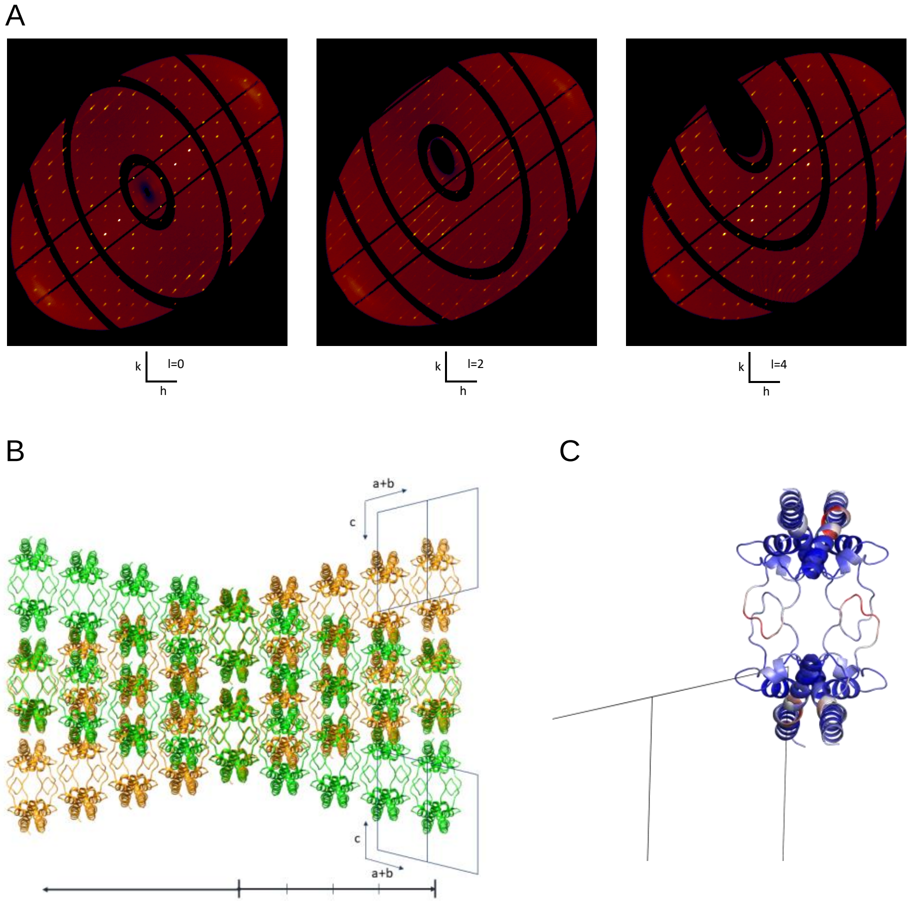


**Figure S3: Twinned CD9$_{EC2}$ crystal.** (A) Reciprocal space reconstructions, hk0, hk2 and hk4. Each slice has Bragg reflections from the two twin lattices. In addition, in hk2 streaks along ***a*+b**** can be clearly observed. All slices hk(l=4n) are ordered. (B) Green and orange structures represent the two twin domains in the crystal. The middle layer is the twin interface with base vectors ***c*** and ***a-b***. From there, the structure can continue in the green or orange direction; the relative shift between the two is 1/4 ***c***. Starting from the middle layer, every fourth layer, the two twin structures exactly overlap. (C) Structure of CD9$_{EC2}$ viewed in the same orientation as (B) and colored by B-factor (spectrum blue-white-red, minimum 20 maximum 100 Å$^2$). A region within the D loop shows high B-factor. It is located at the interface with the next layer, along the ***a+b*** direction.

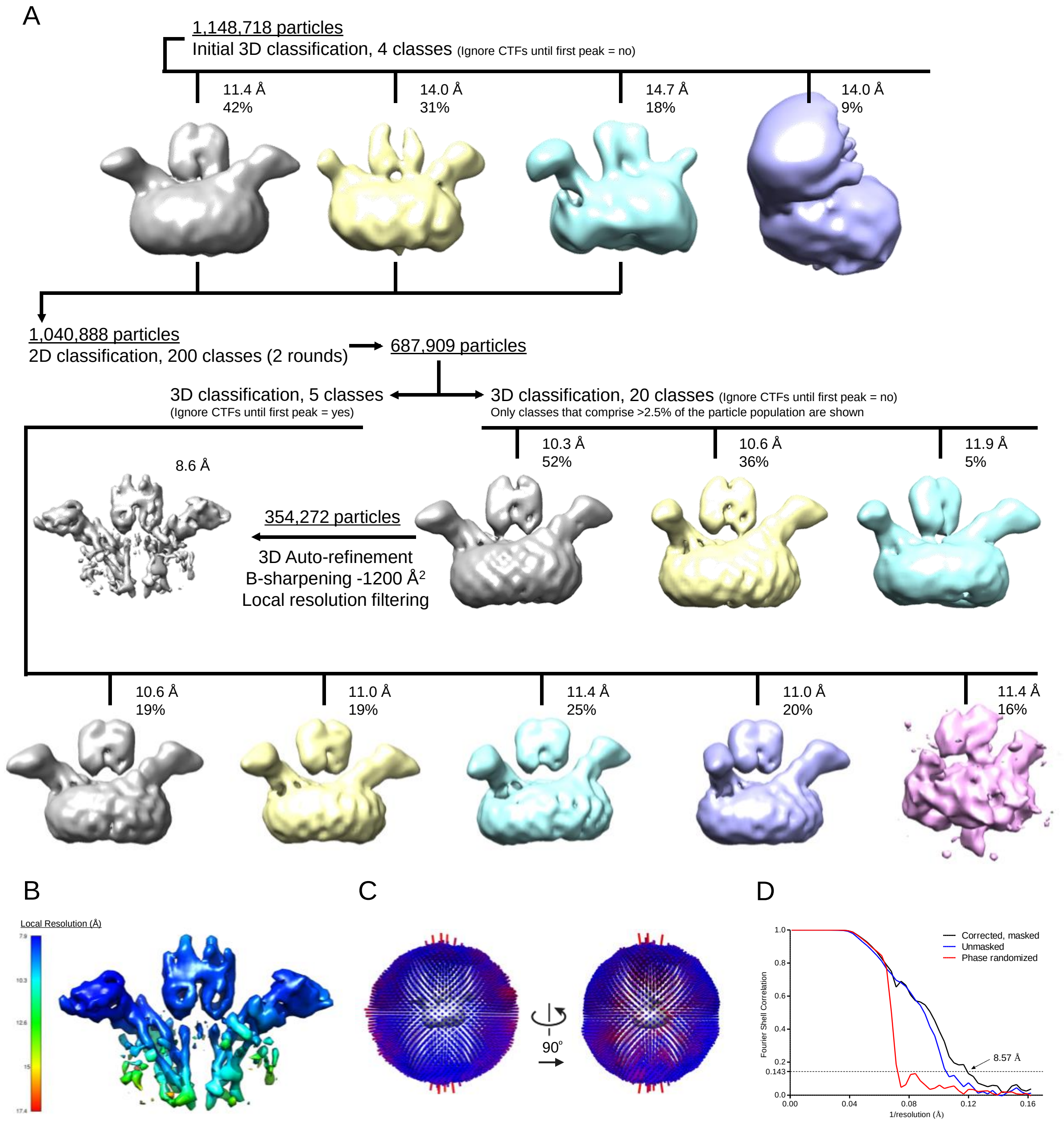


**Fig. S4: Cryo-EM image processing of the EWI-F$_{\Delta Ig1\text{-}5}$ - CD9 - 4C8 dataset.** (A) Image-processing strategy in the Relion pipeline (see experimental procedures). All density maps from a single 3D classification are depicted at the same contour level in the same orientation. (B) Local-resolution estimation of the reconstruction, computed through Relion. (C) Angular distribution of the particles that were used for the reconstruction of the 8.6-Å resolution density map. (D) Fourier-shell correlation plot for gold-standard refined masked (black), unmasked (blue) and high-resolution phase randomized (red) half maps. The FSC=0.143 threshold is shown as a dashed line.

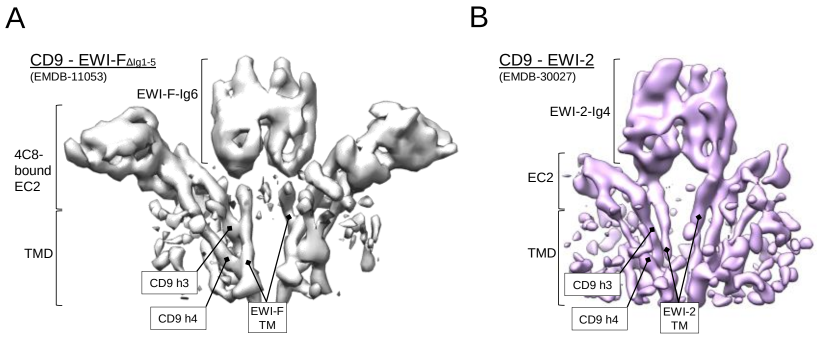


**Figure S5:** (A, B) Cryo-EM density maps of CD9 - EWI-F (from this study) and CD9 - EWI-2 (EMDB-30027) shown in the same orientation parallel to the membrane as a sideview. The density for the Ig1 - Ig3 domains of CD9 - EWI-2 is hidden for clarity. Different protein regions of both maps are annotated
